# The dynamics of stem and crown groups

**DOI:** 10.1101/633008

**Authors:** Graham E. Budd, Richard P. Mann

## Abstract

The fossil record of the origins of major groups is of great interests to many biologists, especially when the fossil record apparently conflicts with timings based on molecular clock estimates. Here we model the diversity of “stem” (basal) and “crown” (modern) members of groups as seen in the fossil record, using a “birth-death model”. Under background conditions, the stem group members must diversify rapidly until the modern crown group emerges, at which point their diversity rapidly collapses, followed shortly by their extinction. Mass extinctions can disturb this pattern to create very diverse stem groups such as the dinosaurs and trilobites. Understanding these null-hypothesis patterns is essential for framing ecological and evolutionary explanations for how major groups originate and subsequently evolve.

---

The fossil record shows many striking patterns [1] that have been addressed by hypotheses such as mass extinctions [2], diversity-dependent diversification [3] and competitive displacement [4, 5]. In particular, the timing and nature of the emergence of modern groups such as animals [6], birds [7, 8] and flowering plants [9] have been of intense interest: here, molecular clock estimates have often been at odds with estimates based solely on the fossil record [9, 10, 11]. Understanding this discrepancy has, however, been hampered by the lack of a model of how both basal (“stem group”) and modern (“crown group”) members [6, 12] of a particular group diversify and go extinct in the fossil record. Quantification of the fossil record, rooted especially in the work of Raup and others[13], allowed explicit models of diversification to be tested, although problems remained about estimation of speciation and extinction rates from the fossil record [14, 15]. Such modelling also failed to take into account the important survivorship and selection biases of the “push of the past” and large clade effects [16], nor the systematic differences that exist between extinct and extant taxa [6, 12]. Any clade can be divided into two components: the last common ancestor of all the living forms and all of its descendants (the “crown group”), and the extinct organisms more closely related to a particular crown group than to any other living group (the “stem group”): together, they make up the “total group” (Fig. 1a). Here we also define the provisional crown group (pCG): the crown group as it would have appeared in the past (e.g., to a Silurian palaeontologist). As basal members of the pCG go extinct through time, the node defining it moves upwards to subtend the next pair of (then) living sister groups, defining a new pCG. When the last member of the stem group goes extinct, the node defining the crown group will be fixed until the present. Nevertheless, the definitive (i.e. modern) crown group will have emerged some time before this, as a subclade of the then pCG, and therefore the definitive crown group will temporarily co-exist with its corresponding stem group. Diversifications can thus be divided into three sequential phases: (1) only the stem group; (2) both stem and crown group; and (3) only the crown group (Fig. 1a).

**Figure 1:**
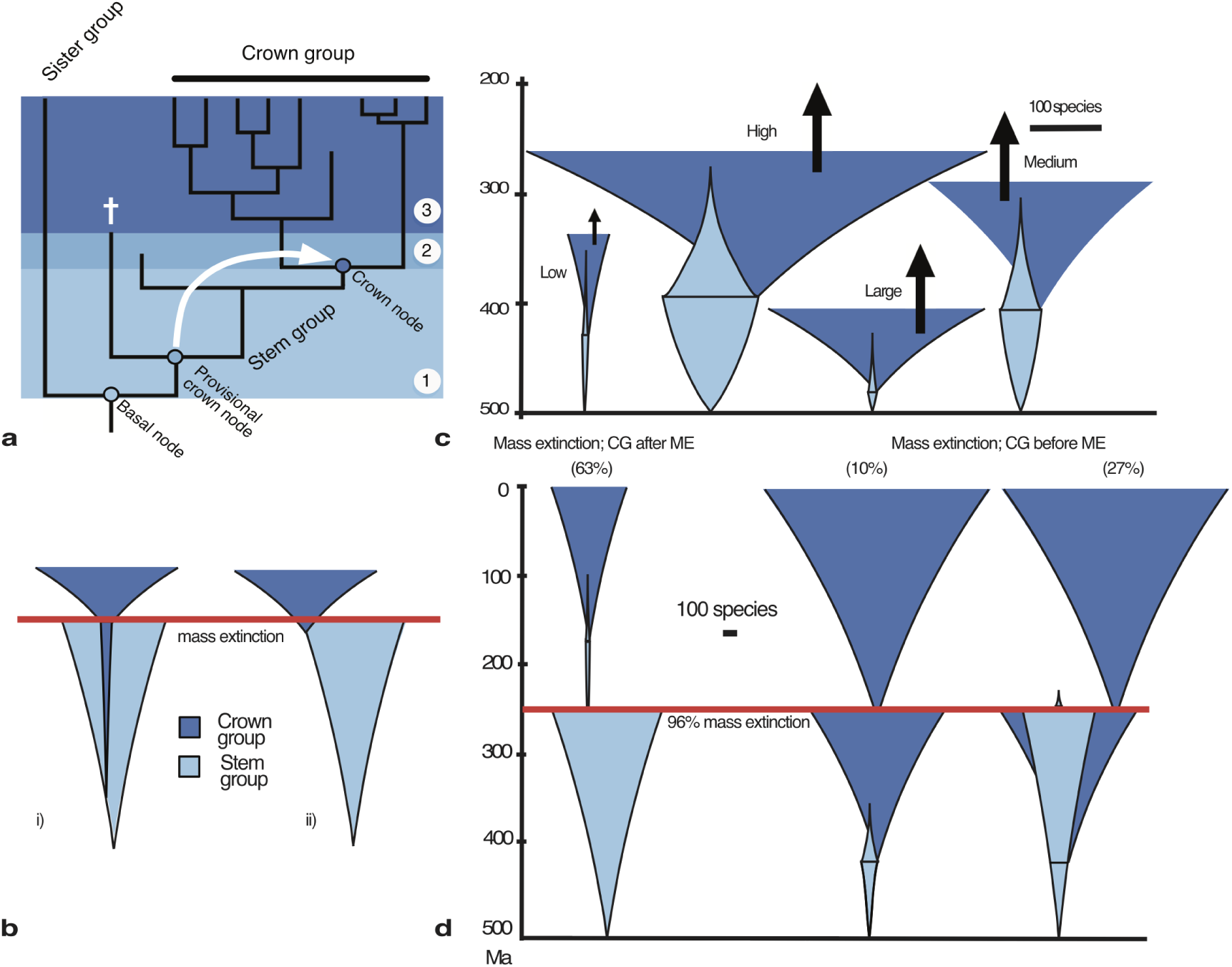
The dynamics of stem and crown groups. **a** Formation of a living crown group is divided into three phases: (1) stem group only; (2) stem and crown group; and (3) crown group only. The definitive crown group is formed as the last stem group member (†) goes extinct, and the previous provisional crown group node moves up to the definitive crown group node (white arrow). **b** Two hypotheses to explain the origin of crown groups: i) the “phylogenetic fuse” [17] where the crown group arises early and is cryptic; ii) origin just before a mass exinction8. **c** To-scale spindle diagrams of typical stem and crown group diversities for low, middle and high turnover rates (extinction rate = 0.1 (with origination rate 0.1143), 0.5 (origination rate 0.5107) and 1 (origination rate 1.0090) per species per million years, each giving c. 10000 species in the present) and a large clade (here five times larger than expected given its diversification parameters18. Note that in all cases (black arrows) the dark blue crown groups continue diversifying to the present. **d** The effect of a large mass extinction (96%) at 250 Ma on the stem and crown groups for a clade with 0.5 background extinction rate that would have been expected to generate 10000 species in the present. The probability of each of the three possible outcomes (crown group after mass extinction; crown group before and stem group dies before; crown group before and stem group reaches mass extinction) are given.

The origins and fates of crown groups are of particular interest as they comprise modern diversity. However, understanding their origins has been hampered by the lack of any analysis of combined stem and crown group dynamics; the plausibility of models to explain the origin of modern diversity (e.g. Fig. 1b) cannot therefore be easily assessed. An absence of crown group taxa after a time of origin based on a molecular clock estimate could be explained by invoking poor fossil preservation [18]. However, observed stem-group diversity potentially offers a mode of assessing this sort of claim if we have a model for relative stem and crown group diversity together. For example: if such a model suggests high crown group diversity when only stem group taxa are observed (e.g. as is the case for birds discussed below), then either the molecular clock estimate for the crown group’s origin is incorrect, or there may be a strong preservation bias between the stem and crown groups that may, on the basis of the model, be further assessed for plausibility.

Here, then, we derive explicit expressions for both stem and crown group diversity and their ratio, using a birth-death model [19]. We first consider how they evolve under homogeneous conditions, and then how mass extinctions perturb the process. In the following, we condition the process on the observation that the clade survives until the present day to form a crown group of at least two species. We take as a model example a total group that emerged 500 Ma, and whose crown group emerged c. 410 Ma; and with diversification parameters speciation *λ* = 0.5107 and extinction *µ* = 0.5 (both per million years) (c.f. fig. 1 of ref. [16]). These numbers would on average generate 10,000 species in the Recent, assuming no mass extinctions. We also consider similarly-sized clades generated with high turnover (*µ* = 1, *λ* = 1.0090) and low turnover (*µ* = 0.1, *λ* = 0.1143), and how unusually large clades [16, 20] (given their diversification parameters) behave. We then introduce mass extinctions [21] of various sizes. In all cases we calculate for 1 million year intervals the expected size of the stem group; the probability that it has gone extinct; the expected proportion of total diversity contained within the stem at any given time (averaged over all clades with these parameters); the probability that the stem contains a certain proportion of diversity, and the expected absolute diversity in the stem (Fig 2). The spindle diagrams of Fig. 1c, d show typical to-scale shapes and sizes of the stem and crown groups under these conditions: note that because of the high stochasticity of the process, a wide range of less-likely outcomes are also possible [16].

**Figure 2:**
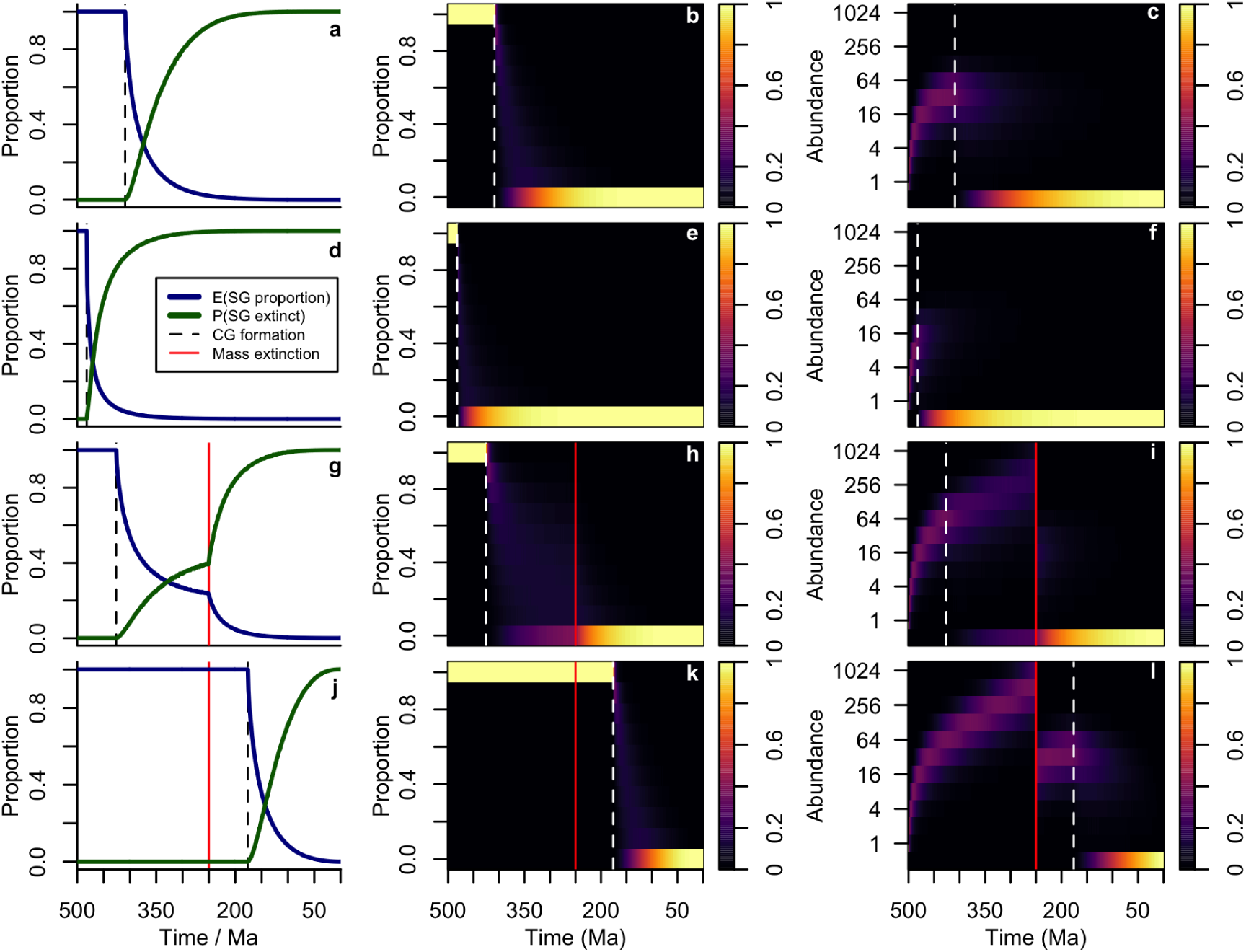
Stem group extinction and diversity. First column: expected total diversity contained in the stem (blue line), probability that stem group has gone extinct (green) line, selected time of origin of crown group (black dashed line) and time of mass extinction (vertical red line); second column; probability heat map of proportion of diversity being contained in the stem group; third column, probability heat map of absolute abundance of stem group through time. In columns 2 and 3, red lines represent a mass extinction and the dashed white line the time of origin of crown group. **a-c**: the “typical” clade of expected size 10000 in the present with speciation rate = 0.5107 and extinction rate = 0.5; **d-f**: a “large clade”, five times larger than expected given its diversification parameters18; **g-i**: the typical clade but with a 96% mass extinction at 250 Ma when the crown group forms before the mass extinction (37% of the time); and **j-l**: as **g-i**, but when the crown group forms after the mass extinction (63% of the time). E(SG proportion): expected proportion of total diversity taken up by stem group; P(SG extinct): proportion of trees in which the stem group has gone extinct. The dashed line represents the time of formation of the crown group (here selected as the mean time from eqn 11, 12 below) and the red line (where present) the time of a mass extinction.

## Results

In our model example, after the emergence of the crown group, the average proportion of diversity contained within the stem group drops exponentially; the stem group makes up half of expected total diversity only c. 15 Myrs after the origin of the crown group. By c. 80 Myrs after, it is expected to make up only 10% of total group diversity. In addition, the probability of the stem group having gone entirely extinct climbs sharply in the same interval: to 50% after 60 Myrs and 75% after 100 Myrs. With high turnover, the stem-group grows faster initially, lasts longer, and generates a larger crown group; however, it too declines extremely quickly once the crown group is established. The converse of all these is true for the low turnover example, except that here too the stem group declines quickly after the crown group origin. These quantitative results illustrate an important insight: crown groups and the stem-group before the crown group emerge rapidly as a necessary condition for survival to the present; and stem groups after the emergence of the crown group cease expansion and rapidly decline as the necessary condition for their extinction. The net effect is that a pCG always diversifies rapidly. The rapidity of the transition from a stem-dominated to a crown-dominated regime is sharpened when the crown group emerges more quickly than expected, as would be the case for an unusually large clade [16, 20]. One would expect the crown group of a clade (such as the arthropods) that ends up perhaps 5 times larger than expected over the same 500 Myrs to emerge at only c. 20 Myrs after the total group (Fig 1b), and for its stem to dwindle very quickly. Provisional crown groups quickly become converted to the definitive crown group, and the crown group node (Fig 1A) then becomes stable. In our example, the definitive crown group forms after an average of 90 Myrs and the stem dies an average of 110 Myrs later, implying that the pCG stops changing from this point and that the identity of the crown group node remains consistent until the present. Such stability explains why the fossil record is dominated by long blocks of time with a very similar biota [2] (e.g. the “Age of Reptiles”). Once a pCG has sufficiently diversified, it becomes very unlikely to go extinct by stochastic variation alone. Such pCGs and their shared characteristics thus become stable features that are disrupted only by the largest mass extinctions.

Long-lived groups have been through one or more mass extinctions. The effect of a very large (96%) mass extinction at 250 Ma on our typical clade is shown in Figs 1d and 2c, d, with the three possibilities of crown group emergence after the mass extinction (63%); crown group emergence before the mass extinction with stem group survival until the mass extinction (27%); or crown group emergence before the mass extinction with the stem group becoming extinct before the mass extinction (10%) being illustrated. The probability of the crown group emerging after a mass extinction is only high when the absolute number of surviving species is small (Fig 3b, c). For our example clade, the tipping point mass extinction size (when the crown group becomes more likely to emerge after than before the extinction) is as much as 93%. Well-established (i.e. relatively long-lived) provisional crown groups, like crown groups themselves, are also robust even to large mass extinctions; the loss of famous clades such as trilobites seems to be the result of successive biotic crises, not a single event [22].

**Figure 3:**
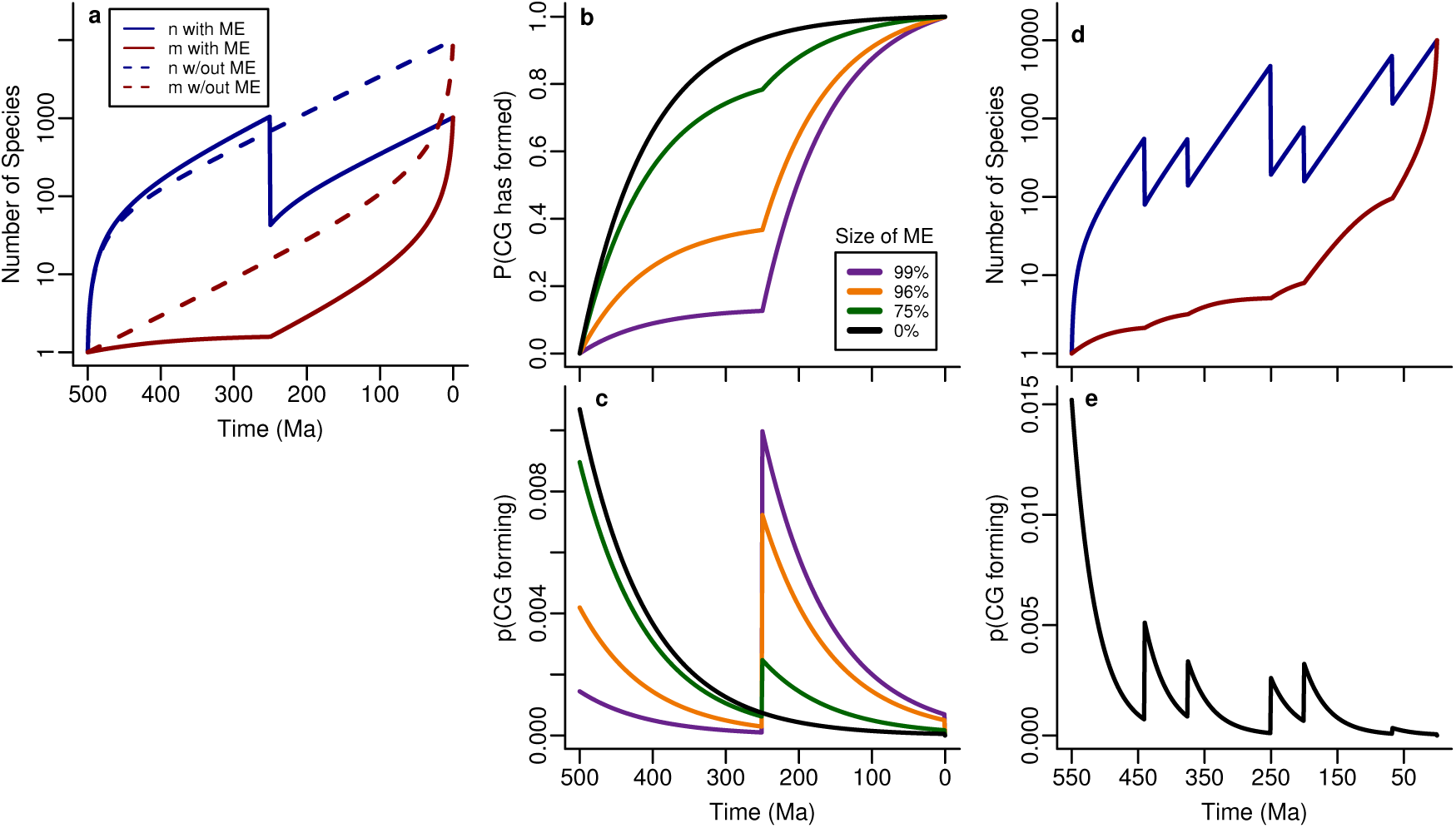
The effects of single and multiple mass extinctions. **a** Species (blue) and lineage (red) plots for a typical diversification that would normally generate 10,000 species in the present over 500 Myrs with background extinction rate 0.5 and speciation rate c. 0.5107 (dotted lines) but is affected by a 96% mass extinction at 250 Ma (note slightly higher initial push of the past18). **b, c** Cumulative and non-cumulative probability distribution curves for probability that crown group has formed for mass extinctions of various magnitudes; note sharp increase in all cases directly after the mass extinction, and that only the largest mass extinctions have a significant effect. **d, e** The effect over Phanerozoic time (c. 550 Myrs) of the canonical “big five” mass extinctions [23] on a typical clade originating at the beginning of the Cambrian on **d** species and lineage counts and **e** probability of crown group forming.

In Fig. 3b, c we show the fates of both species and lineages counts through time as they pass through a mass extinction of varying size (for that particular clade) and the probability of the emergence of the crown group against time. The bifurcating effect of mass extinctions on crown group origins implies that the least likely time for a crown group to emerge is just before a mass extinction (Fig. 3c). For example, if crown Aves really evolved at around 70 Ma8 and *Vegavis* [24] is an anseriform, then at least five crown group bird lineages independently crossed the K/Pg boundary [24] and at the same time had an origin very close to it in time. Our modelling shows this to be an unlikely possibility (c.f. ref. [25]), and that the controversial “explosive” post-Cretaceous model [10] of crown group birds is more likely.

It is also possible to calculate directly the effect of multiple mass extinctions [21], such as the canonical “big five” of the Phanerozoic [23]. For a set of such extinctions across Phanerozoic time (Fig. 3 d, e) it can be seen that the probability of the crown group of a Cambrian total group emerging is sharply higher in periods of time just after mass extinctions, apart from the continued high likelihood of a Cambrian origin. The suggestive resemblance of Fig 3e to plots of Phanerozoic high-level taxonomic origination rates [3, 14] may not be coincidental, with higher level taxa such as families that appear to preferentially appear after mass extinctions corresponding partly to crown groups.

## Discussion

Our results reveal much about the dynamics of stem and crown groups and have an important bearing on how we view the fossil record. In particular, as the crown group forms, the stem group quickly collapses into first obscurity and then extinction unless the total group was affected by a very large mass extinction. Indeed, this rapid phase transition from stage 1 to stage 3 of the diversification process might be mistaken for a mass extinction itself, such as has been suggested for the end of the Ediacaran period [26]. Qualitative models [17] wherein crown groups form “silently” and remain as low diversity components of the background while the stem group diversifies (Fig. 1b) can be seen to be unlikely under the model; rather, the reverse is always true. This is particularly important in clades where molecular clocks suggest a deep origin of the crown group during an interval of time when the stem group is seen to be diversifying in the fossil record, with classic examples being animals, angiosperms, birds and mammals [27, 9, 28, 18, 11] (Fig. 1b). In common with other recent analyses [9, 16, 27], this suggests that systematic errors in molecular clock estimates, especially for large clades, are creating illusory deep roots that are profoundly at odds with both modelling of diversification and observed patterns in the fossil record.

The famous debates in the 1970s in the pages of Paleobiology and elsewhere [29] were essentially between viewing the fossil record as an archive of discernable evolutionary processes (such as diversity-dependent diversification [3]) or as the result of inevitable stochastic patterns that could be captured with simple statistical models [30]. We have previously extended the latter view by modelling heterogeneity that results from survivorship and selection bias [16]. Here we show that other types of heterogeneity in the record such as the rapid takeover of diversity by the crown group and preferential crown group emergence after large mass extinctions also emerge from imposing retrospective structure on a homogeneous process and should therefore also be part of any null hypothesis. A homogeneous model is not of course, intended to capture the many true heterogeneities that exist in the evolutionary process: it is a model, not a description. Nevertheless, it shows both that strong patterns of heterogeneity emerge even under homogeneous background conditions when simple conditions such as survival to the present are imposed, and also that hypotheses that strongly transgress against the general predictions of the model should not simply be accepted without scrutiny, especially when such transgressions are themselves generalised (e.g. the “phylogenetic fuse” [17] concept).

## Mathematical methods

The fundamental mathematical results concerning the distribution of species and lineage numbers in a birth-death process are given by Nee et al. [19], and have been much elaborated and applied subsequently, especially for the reconstructed process (i.e. models for lineages leading to extant species) [21, 31, 32]. Here we use those results to derive properties of the stem and crown groups, both in homogeneous processes and those experiencing mass extinctions that occur at singular points in time, with references to related approaches for the reconstructed process where appropriate. Our notation follows that used in Budd and Mann [16], but we strive to make clear the connection to the notation of Nee et al. [19] and others.

### Glossary of mathematical terminology

- *λ*: the speciation rate of a birth-death process (BDP), assumed to be constant.
- *µ*: the extinction rate of a BDP, assumed to be constant.
- *T*: the time from the origin of a BDP to the present day. Equivalently the age of the total group.
- *n*_*t,t*′_: the number of species alive at time *t*′ in a BDP that originated at time *t*.
- *m*_*t,t*′_: the number of species alive at time *t* with descendants at time *t*^′^, in a BDP that originated at time zero.
- *a*_*t,t*′_: parameter controlling the distribution of abundance such that (1 − *a*_*t,t*′_) is the “succes” parameter in the geometric distribution of *n*_*t,t*′_ (see equation 1).
- *S*_*t,t*′_: the probability that a BDP originating at time *t* has any descendants at time *t*′.
- *f*_*i*_: the fraction of species that survive mass extinction event *i*.
- 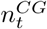: the number of species at time *t* in the crown group of a BDP that originated at time zero.
- 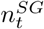: the number of species at time *t* in the stem group of a BDP that originated at time zero.

### The birth-death process

The number of living species, *n*_*t,t*′_ at time *t*^′^ in a process that originates at time *t*, conditioned on the survival of the process to *t*′ is geometrically distributed with success parameter 1 − *a*_*t,t*′_:

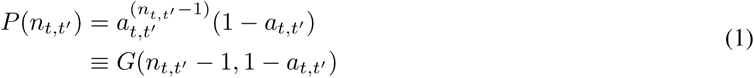

The parameter *a*_*t,t*′_ is analogous to the *u*_*t*_ of Nee et al. [19], and also corresponds to the lower tail distribution of the coalescent point process *P* (*H* < *T*) described by Lambert and Stadler [21], where *H* is the coalescence time and *T* is as above. In a homogeneous process without mass extinctions, *a*_(_*t, t*′) is a function of the background speciation rate *λ* and extinction rate *µ*:

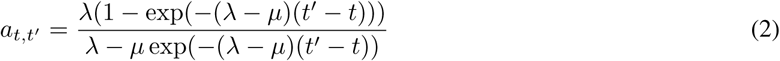

Nee et al.[19] showed that the number of species alive at time *t* with descendants at time *t*′, *m*_*t,t*′_ is also geometrically distributed (note Nee et al. [19] use the notation *η* rather than m):

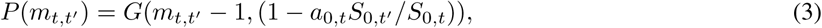

where *S*_*t,t*′_ is the probability for a birth-death process originating with one species at time *t* to survive to time *t*′ (analogous to the *P* (*t, t*′) of Nee et al.[19]). In a homogeneous process this survival probability is given as:

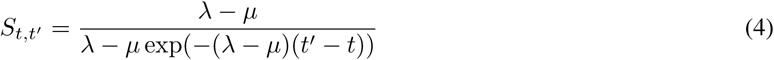

### Mass extinctions

As shown by Nee et al. [19], the distributions of *n*_*t,t*′_ and *m*_*t,t*′_ in terms of *a*_*t,t*′_ and *S*_*t,t*′_ continue to hold when the process is not homogeneous, for example when mass extinctions occur. However, the forms of *a*_*t,t*′_ and *S*_*t,t*′_ must change to account for this heterogeneity. The parameter controlling species diversity, *a*_*t,t*′_, can be given more generally in terms of time dependent *λ*(*s*) and *µ*(*s*) as:

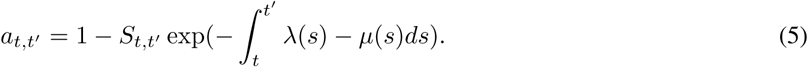

In the case of a set of mass extinctions occurring instantaneously at times between *t* and *t*′, with survival fractions *f*_1_, *f*_2_, …, *f*_*N*_, in an otherwise homogeneous process, this reduces to:

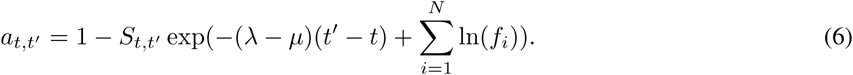

What remains is to determine the survival probability *S*_*t,t*′_ of the process through the mass extinctions. First we consider survival through one mass extinction, and then identify a recursive process for calculating the survival probability through multiple mass extinctions. Consider a birth death process starting with one species at time *t*. If each species has a instantaneous and independent probability *f* to survive a mass extinction at time *t*_*m*_, what is the probability that the process has descendants at time *t*′, beyond the mass extinction?

Firstly, the distribution of the number of descendants it has at time *t*_*m*_, from equation 1, is:

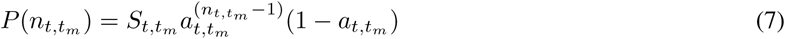

The probability that any one of these descendants at time *t*_*m*_ will leave a descendent at time *t*′ is given by the product of the (independent) probability that it survives the mass extinction (*f*) and the probability that its lineage survives another time *t*′ − *t*_*m*_. Therefore the probability that at least one will survive is:

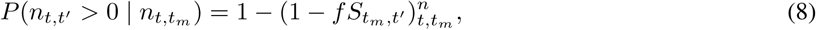

To obtain the probability *S*_*t,t*′_ that any species at time *t* will have some descendants at time *t*′ we must marginalize over the unknown value of 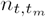:

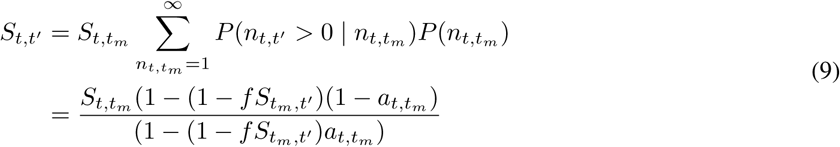

In the case of one mass extinction, 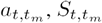 and 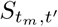 are all given by equations 2 and 4, since these periods contain no further heterogeneities. However, in the case of multiple mass extinctions, equation 9 can be applied recursively along with equation 6 to obtain these values, until all mass extinctions have been incorporated.

Using these adapted expressions for *a*_*t,t*′_ and *S*_*t,t*′_, the geometric distributions specified in equations 1 and 3 continue to hold19. We note that Lambert and Stadler [21] considered the reconstructed evolutionary process as a coalescent point process (CPP), and concisely derived a full distribution of trees through an arbitrary set of mass extinctions, showing that the process remains a CPP. This prior result thus provides an alternative derivation for our results concerning the number of lineages that survive to the present, *m*_*t,T*_, as these constitute the reconstructed phylogeny. However, the reconstructed process by definition excludes the extinct stem lineages.

### When does the crown group form?

The probability that the crown group has formed by time *t* is the probability that the number of species alive at time *t* that have descendants in the present, *m*_*t,T*_ is greater than one. We take a crown group here to consist of at least two species, so that both the crown and stem group can be defined. Since *m*_*T,T*_ > 1 (i.e. we observe that the crown group does form before the present), this implies:

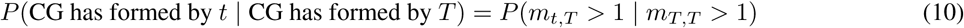

Using equation 3 we have:

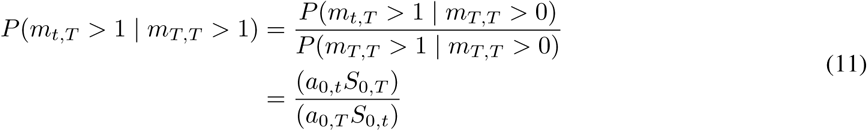

We can further condition on a known limit to the time of the crown group. For example, if we wish to consider cases where the crown group emerges before the time of a mass extinction, *t*_*m*_, the following the above gives an updated expression:

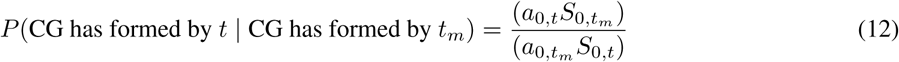

This result can also straightforwardly be derived by the CPP approach of Lambert and Stadler [21] since the crown group node subtends all extant species of a clade.

### Dynamics of stem and crown group abundance

In this section we derive the new mathematics required to model stem and crown group abundance together. We assume that the process starts at time *t* = 0 and that the time at which the crown group emerges, *t*_*c*_ is known, and derive expressions for abundance in the stem and crown groups conditioned on this.

### Before the crown group forms

Before the crown group has formed, all diversity is stem diversity. We can calculate the distribution of this diversity, 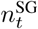, by conditioning on *m*_*t,T*_ = 1, i.e. that the crown group has not yet formed, revealing a negative binomial distribution:

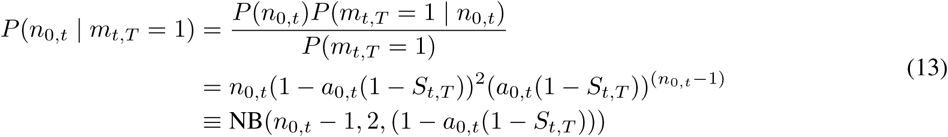

### After the crown group forms

#### The crown group

The crown group is defined by a speciation at time *t*_*c*_, creating exactly two species that will give rise to descendants in the present. This implies two independent BD processes that will survive to time *T*, starting at time *t*_*c*_, thus implying for each that 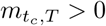. This implies [19] that the abundance for each of these processes is the sum of two non-identically distributed geometric random variables, *X* and *Y*, with:

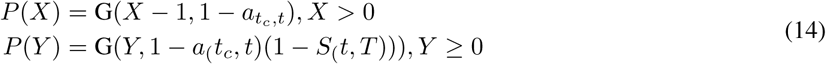

Since the crown group consists of two such independent BD processes, we can use the fact that the sum of two i.i.d. geometric random variables follows a negative binomial distribution to express the crown group diversity at time *t*:

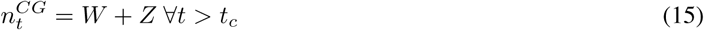

with

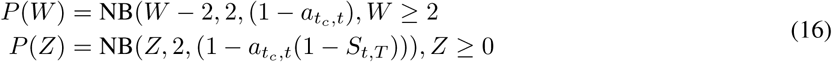

#### The stem group

Let 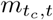 be the number of species just before the crown group speciation event that give rise to descendants at time *t*. We know that 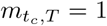, since one species will form the base of the crown group, while the others will give rise to the stem, which is extinct at time *T*. Conditioning on this, we have:

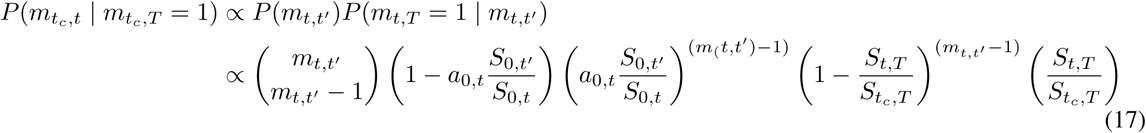

With correct normalisation this reveals a negative binomial distribution:

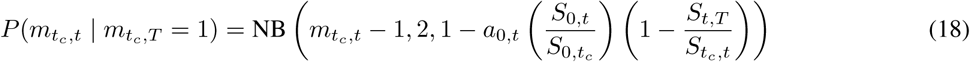

We can identify the quantity 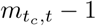 as the number of species at time *t*_*c*_ prior to the crown group speciation that have descendants in the stem at time *t*, since one species at time *t*_*c*_ only has descendants in the crown group. The total stem diversity at time *t* is therefore the sum of 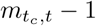 independent BD processes that are conditioned on surviving for a duration *t* − *t*_*c*_ but also on becoming extinct before time *T* (by definition the stem is extinct at time *T*). First we consider the dynamics of one such BD process. From equation 1, conditioning only on survival to time *t* from a start of *t*_*c*_, we have:

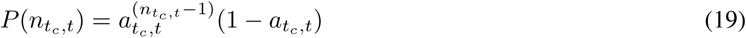

Now we also condition on the process not surviving to time *T*:

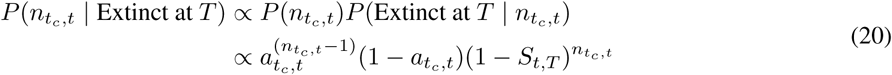

With normalisation this reveals another geometric distribution:

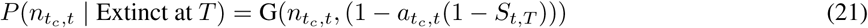

We can now see the total stem abundance at time *t* as the sum of *m*_(_*t*_*c*_, *t*) − 1 i.i.d. geometrically-distributed random variables, where *m*_(_*t*_*c*_, *t*) − 1 is itself composed of the sum of two geometrically distributed random variables (by virtue of its own negative binomial distribution). Marginalising over possible values of 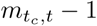, this gives a full marginal distribution for the stem diversity 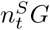 for times *t* > *t*_*c*_:

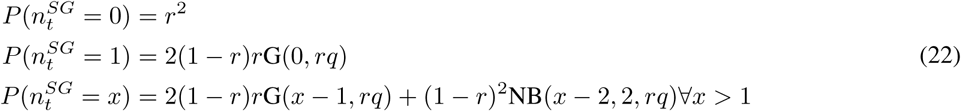

where 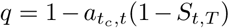 and 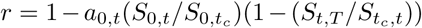, and the geometric and negative binomial mass functions G and NB are defined as above. A direct corollary of this result is that the probability that the stem is extinct at time *t* is *r*^2^.

